# Local modulation of sleep slow waves depends on timing between auditory stimuli

**DOI:** 10.1101/2025.03.05.641406

**Authors:** Sven Leach, Sara Fattinger, Elena Krugliakova, Jelena Skorucak, Georgia Sousouri, Sophia Snipes, Selina Schühle, Maria Laura Ferster, Giulia Da Poian, Walter Karlen, Reto Huber

## Abstract

Conflicting evidence exists regarding the role of the targeted slow-wave phase in determining the direction and spatial specificity of slow-wave activity (SWA) modulation via phase-targeted auditory stimulation (PTAS) during sleep. To reconcile these discrepancies, we re-analyzed high-density electroencephalography (hd-EEG) data from previous studies, focusing on SWA responses to auditory stimuli presented with varying inter-stimulus intervals (ISIs). Our analysis reveals that ISI is a primary determinant of PTAS-induced SWA modulation, exceeding the influence of targeted phase alone. Specifically, auditory stimulation with longer ISIs evoked a global increase in SWA, consistent with a stereotypical auditory-evoked K-complex (KC), independent of targeted phase. Conversely, longer stimulus trains with rapid successive stimulus presentation resulted in spatially localized, phase-dependent SWA modulation, with up-PTAS enhancing and down-PTAS reducing SWA locally around the targeted area. This distinction resolves inconsistencies in prior PTAS studies by demonstrating that phase alone in insufficient in predicting slow-wave responses. Rather, it was the ISI which determined whether PTAS resulted in a global, KC-mediated response or a local, phase-specific modulation of SWA. Consequently, our findings refine the mechanistic understanding of PTAS, suggesting that ISI regulates the engagement of distinct neural circuits and thereby potentially enables the targeted manipulation of specific slow-wave subtypes and their associated functions.

## Introduction

Neural oscillations—rhythmic brain activity patterns evident in the electroencephalogram (EEG)—underlie various aspects of human behavior. Their study is primarily motivated by the desire to understand the computational principles governing core neural processes (Cohen, 2017). Neuromodulation techniques seek to experimentally manipulate such oscillations to study concurrent behavioral consequences, offering a promising approach to establishing causal relationships between the two (Herrmann et al., 2016). Neuromodulation during sleep has placed particular emphasis on slow waves (0.5–4 Hz) and sleep spindles (12–16 Hz), two prominent EEG oscillations during non-rapid eye movement (NREM) sleep which are thought vital for sustaining a wide range of physiological functions (Fernandez & Lüthi, 2020; Léger et al., 2018).

Given its simplicity and non-invasiveness, the modulation of slow waves with non-arousing auditory stimuli has received substantial attention over the last decade. At its core, phase-targeted auditory stimulation (PTAS), also known as phase-locked (PLAS) or closed-loop auditory stimulation (CLAS), involves the presentation of auditory stimuli locked to a specific phase of ongoing slow waves (Ngo et al., 2013). Using PTAS, several studies have suggested causal relationships between slow waves and memory consolidation (Clark et al., 2024; Diep et al., 2020; Leminen et al., 2017; Moreira et al., 2021; Ngo et al., 2013; Ong et al., 2016, 2018; Papalambros et al., 2017; Prehn-Kristensen et al., 2020; Salfi et al., 2025), immune functions (Besedovsky et al., 2017), cardiovascular health (Grimaldi et al., 2019; Huwiler et al., 2023, 2024), learning capacity (Fattinger et al., 2017), and general sleep-related recovery processes (Krugliakova et al., 2022).

The central premise of a *conceptual framework* that describes how PTAS is thought to modulate neural oscillations during sleep is that the phase of targeted slow waves directly relates to neuronal firing (or silence) of cortical neurons (Nir et al., 2011; Steriade et al., 1993; Vyazovskiy et al., 2009). An auditory stimulus delivered during the up-phase of slow waves (up-PTAS), a depolarized state as active as during wake, is thought to contribute to the existing neuronal activity and therefore further enhance neuronal synchrony. Conversely, targeting the down-phase (down-PTAS), a hyperpolarized state where neurons are completely silenced for a few hundred milliseconds, is proposed to disrupt neural synchrony by perturbing this silenced state (Bellesi et al., 2014). The second central aspect is the locality of PTAS effects. Most slow waves do not occur synchronously across the scalp but originate in one location and travel across the cortex (Massimini et al., 2004; Sousouri et al., 2022). Phase-locked stimuli are delivered based on the EEG signal from a single detection electrode. As a result, systematic phase targeting occurs primarily near the detection electrode, with precision diminishing as the distance from this electrode increases. This explains how an auditory stimulus, despite reaching a large portion of the cortex, could interact systematically with only a circumscribed region.

The current landscape of data, however, partly conflicts with this framework: (1) While up-PTAS has consistently resulted in enhancements of slow waves, this effect has never been demonstrated to be local (Huwiler et al., 2022; Krugliakova et al., 2022); (2) while down-PTAS should exclusively result in the local suppression of slow waves (Fattinger et al., 2017; Moreira et al., 2021; Ngo et al., 2013), also global enhancements thereof have been observed (Huwiler et al., 2022; Leach et al., 2024). These incongruent results seriously question the current conceptual framework of PTAS and therewith the derived conclusions from studies that applied PTAS.

This work aims to reconcile these conflicting results by introducing a critical new ingredient to the framework: *inter-stimulus intervals (ISIs)*. In our previous work, we observed that PTAS induces a K-complex-like (KC-like) response, regardless of the targeted phase, particularly when stimuli were spaced apart (Leach et al., 2024). Since KC induction habituates with repeated stimulus presentation, potentially due to inherent refractory periods (Colrain, 2005), a stimulation protocol which presents stimuli irregularly and infrequently would facilitate KC induction by periodically interrupting habituation processes or resetting refractory periods. Indeed, the only study which reported a local modulation of slow-wave activity (SWA; spectral power between 1 and 4 Hz) presented stimuli continuously throughout sleep (Fattinger et al., 2017), facilitating stimulus habituation and suppressing KC induction. In contrast, studies which reported global SWA increases restricted stimulus presentation to periodically occurring ON windows with durations of a few seconds (Huwiler et al., 2022; Krugliakova et al., 2022; Leach et al., 2024), allowing the regular interruption of stimulus habituation and facilitating KC induction. In these studies, the largest increase in SWA was observed with the first stimulus within an ON window, supporting the idea that habituation was lost during intermittent stimulation breaks. Consequently, we hypothesized that a local, *phase-specific response* could only occur when stimuli bypass KC induction, for instance, when being presented regularly and in rapid succession. To test this, we reanalyzed two previously published datasets (see Methods) in which participants underwent at least two separate laboratory nights: one with up- or down-PTAS, and another without any stimulation (SHAM). We indeed found that PTAS induces local, phase-dependent modulations of slow waves only when stimuli were presented in rapid succession within trains.

## Results

To test whether PTAS responses depend on the *timing between stimuli*, we reanalyzed high-density EEG (hd-EEG) data recorded from 128 channels in ten participants during a night with and without up-PTAS, and from fourteen participants during a night with and without down-PTAS (see Methods). In these studies, stimuli (50 ms pink noise) were presented during ON windows (6 s and 16 s for up- and down-PTAS, respectively), allowing stimulation, which alternated with OFF windows (6 s and 8s), withholding stimulation. Phase-targeting was achieved using EEG from a detection electrode near FP2 (up-PTAS) or C3 (down-PTAS). We previously observed that spectral responses following PTAS closely resembled those of auditory evoked KCs when stimuli followed a long stimulation-free period (Leach et al., 2024). Analogously, we here compare the event-related spectral perturbation (ERSP) following stimuli which were preceded by either long (*≥* 5 seconds) or short (≤ 1 second) stimulation-free periods after both up- and down-PTAS.

### K-complex response

Replicating previous findings (Leach et al., 2024), both up- and down-PTAS resulted in a stereotypical KC-response profile when considering isolated stimuli with long prior stimulation-free periods (**Fig. 1A** & **Fig. 2A**). This response was characterized by a global enhancement of SWA in an early, pre-defined time window (1 – 4 Hz; 300 – 700 ms) following both up-PTAS (max. *g* = 1.17; mean g = 0.66; cluster of 100 channels; see Methods) and down-PTAS (max. *g* = 1.66; mean *g* = 1.10; cluster of 110 channels), compared to SHAM (p <. 05, cluster corrected; **Fig. 1A**). As anticipated for KC responses, SWA increases were accompanied by a global enhancement in theta activity (4 – 8 Hz; 300 – 700 ms) following both up-PTAS (max. *g* = 1.12; mean *g* = 0.81; cluster of 103 channels) and down-PTAS (max. *g* = 1.65; mean *g* = 1.29; cluster of 110 channels), compared to SHAM (p < .05, cluster corrected; **Supplementary Fig. 3**), as well as sigma activity (12 – 16 Hz; 0.9 – 1.5 s) following both up-PTAS (max. g = 0.90; mean g = 0.66; cluster of 106 channels) and down-PTAS (max. *g* = 0.80; mean *g* = 0.59; cluster of 110 channels), compared to SHAM (p < .05, cluster corrected; **Supplementary Fig. 4A**).

**Fig. 1:**
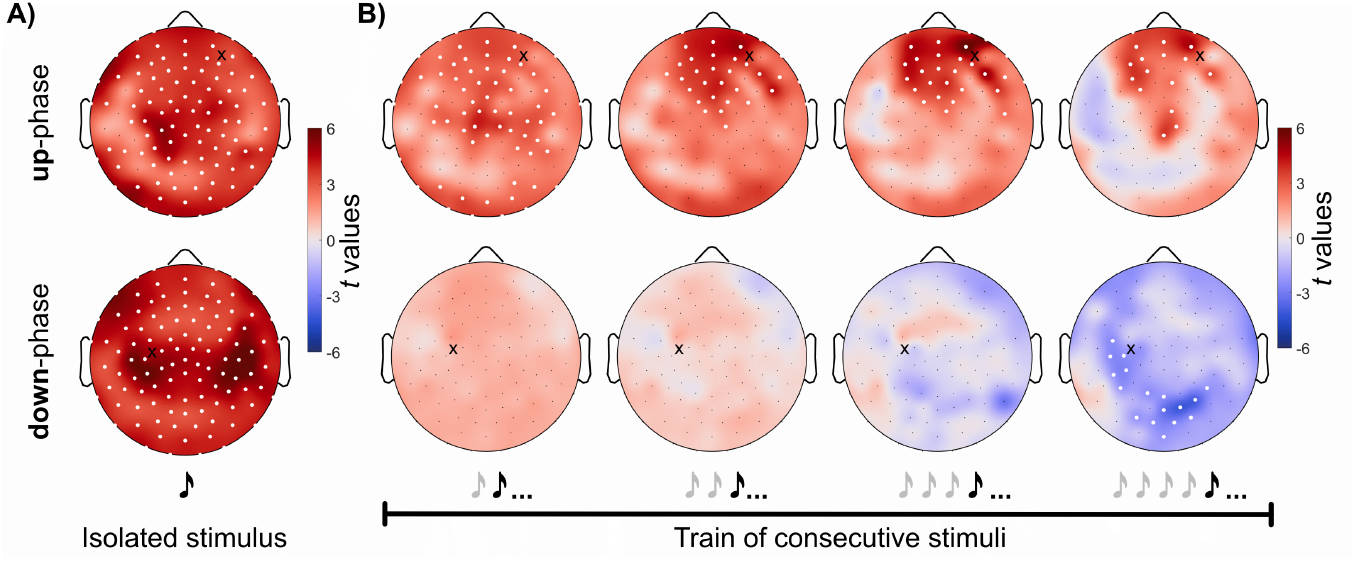
Slow-wave activity response to isolated stimuli and those presented consecutively within trains. **(A)** Topography of absolute slow-wave activity (SWA; 1 – 4 Hz; 300 – 700 ms following stimulus onset) in response to isolated stimuli, that is, stimuli following a stimulation-free period of *≥* 5 s. Responses are displayed either for stimuli targeting the up-phase (top) or down-phase (bottom) of slow waves. **(B)** Topography of absolute SWA (1.2 – 2.6 s) in response to stimuli within a train of consecutive stimuli, that is, stimuli with an inter-stimulus interval (ISI) ≤ 1 s. The average response to a stimulus and all subsequent stimuli within the train rather than the response to a stimulus at a particular train position is shown (indicated by the ellipsis, the three dots; see **Supplementary Fig. 2A** for trial numbers). **Symbols:** The black cross indicates the detection electrode. White dots represent electrodes with significant differences between conditions (STIM vs. SHAM, paired *t*-test, cluster corrected). **Take-away:** The local, phase-specific response becomes more local over time and occurs faster when targeting the up-compared to the down-phase of slow waves, potentially as both the K-complex and phase-specific response following up-PTAS share the same direction of effect.

**Fig. 2:**
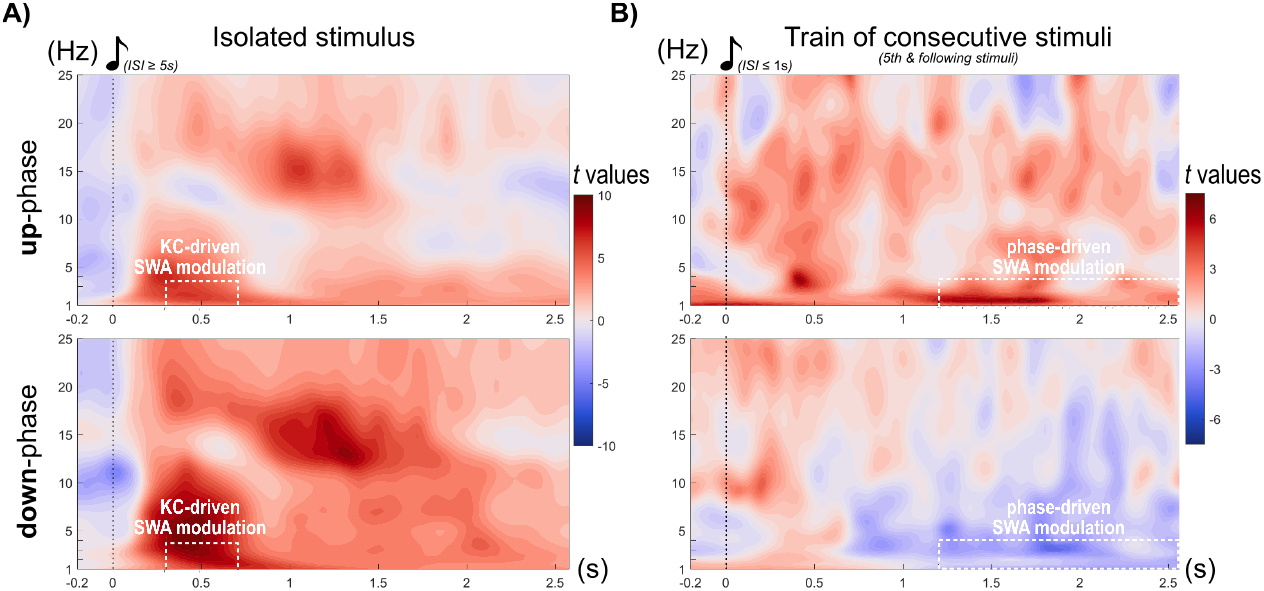
Time-frequency response to isolated stimuli and those within later train positions. Normalized (see methods) event-related spectral perturbation (ERSP) following stimuli targeting either the up- (top) or down-phase (bottom) of slow waves. **(A)** The ERSP of the average of all channels following isolated stimuli, that is, stimuli following a stimulation-free period of *≥* 5 s, is shown. K-Complex (KC) characteristics, encompassing increases in SWA, theta, and sigma activity, are clearly visible after both up- and down-phase stimulation. The topography of the global slow-wave activity (SWA) response (300 – 700 ms; 1 – 4 Hz, indicated by the white dashed box) is depicted in **Fig. 1A. (B)** ERSP in response to stimuli following a train of stimuli with an inter-stimulus interval (ISI) ≤ 1 s. The response to the 5th and all subsequent stimuli within a given train was averaged. The topography of the localized SWA response (1.2 – 2.6 s; 1 – 4 Hz, indicated by the white dashed box) is depicted in **Fig. 1B**. This later time period was chosen to minimize potential interference from KC responses as observed in **Fig. 1A**. To account for the locality of this response, the ERSP depicted here represents an average across channels showing a significant difference in SWA between conditions in **Fig. 1B. Takeaway:** Isolated stimuli elicit a global, stereotypical KC response, regardless of the targeted slow-wave phase. In contrast, following a train of stimuli presented in rapid succession, a local, phase-specific response occurs: targeting the up-phase of slow waves locally enhances, while targeting down-phase of slow waves locally decreases SWA.

### Phase-specific response

Conversely, when stimuli were presented in rapid succession, we observed diverging and localized PTAS effects which depended on the targeted phase after the fifth stimulus in a train of such stimuli (**Fig. 1B** & **Fig. 2B**). Topographic analyses of SWA responses during a later time window (1 – 4 Hz; 1.2 – 2.6 s; minimizing potential interference from co-occurring KC responses) revealed locally enhanced SWA following up-PTAS in a cluster of 19 frontal channels (max. *g* = 0.98; mean *g* = 0.33), and locally reduced SWA following down-PTAS in a cluster of 19 centro-parietal channels (min. *g* = -0.34; mean *g* = -0.22), compared to SHAM (p <. 05, cluster corrected; **Fig. 1B**).

Notably, following up-PTAS, an increase in SWA was already detectable after the second tone (max. *g* = 0.75; mean *g* = 0.28; cluster of 68 channels) and became progressively more localized with each subsequent stimulus (*3*^*rd*^ *stimulus*: cluster of 34 channels; max. *g* = 0.96; mean *g* = 0.33; *4*_*th*_ *stimulus*: cluster of 30 channels; max. *g* = 0.90; mean *g* = 0.31). In contrast, with down-PTAS, the phase-dependent response was first evident after the fifth stimulus. Consistent with the reported refractory period of thalamocortical cells underlying spindle generation (Fernandez & Lüthi, 2020), this global increase in sigma activity was observed specifically for stimuli that followed long stimulation-free periods (all p > .05, cluster corrected; **Supplementary Fig. 4B**).

## Discussion

The precision with which auditory stimuli align with the targeted phase of slow waves has traditionally been considered the single most important factor in determining the outcome of PTAS (Navarrete et al., 2020; Ngo et al., 2013). However, conflicting results in PTAS studies already suggest that the targeted phase is insufficient to fully explain its outcomes. This work introduces another factor which appears to be at least equally important for predicting PTAS outcomes: *inter-stimulus intervals*.

Specifically, our findings indicate a dual-response to PTAS, where stimuli can both (1) lead to a global enhancement of SWA, irrespective of the targeted phase, likely through the induction of KCs and (2) locally modulate SWA, with up- and down-PTAS either increasing or reducing SWA, respectively. Importantly, the nature of these responses was determined by the length of the stimulation-free period prior to a given stimulus, with longer intervals promoting a global enhancement of SWA and shorter intervals allowing the local modulation of SWA.

As early as the 1930s, researchers observed that sensory stimuli, particularly auditory stimuli (Davis et al., 1939), could evoke KCs during sleep (Loomis et al., 1938). The term “K-complex” itself likely derives from the “knock” stimuli used in those early studies (Halász, 2016). It is therefore only logical that auditory stimuli, even when timed to specific slow-wave phases, can still evoke KCs. The observation of either a global increase in SWA following PTAS (Huwiler et al., 2022; Krugliakova et al., 2022), or a stereotypical KC spectral response profile, encompassing increases in SWA, theta, and sigma activity (Krugliakova et al., 2020, 2022; Leach et al., 2024; Ong et al., 2016, 2018), would therefore strongly implicate KCs as at least a partial contributor to the observed modulation of SWA in these studies.

Conversely, rapid stimulus presentation has been shown to diminish the likelihood of eliciting subsequent KCs (Davis et al., 1939; Roth et al., 1956). Consistent with this, here, both up- and down-PTAS were capable of modulating SWA locally when stimuli were presented in rapid succession within trains, thereby essentially bypassing KCs, with up-PTAS increasing and down-PTAS reducing SWA, respectively. Thus, these findings serve as first evidence suggesting that a phase-specific response, spatially confined to areas around and posterior to the detection electrode, likely due to slow-wave traveling (Massimini et al., 2004; Sousouri et al., 2022), unfolds once stimuli are presented consecutively within trains. Our findings indicate weaker phase-specific SWA modulations compared to KC responses, as reflected in the smaller effect sizes for both up-PTAS (max. *g* = 1. 17 vs. 0.98) and down-PTAS (max. *g* = 1. 66 vs. -0.34).

Crucially, this insight reconciles conflicting results from prior studies, where down-PTAS was reported to either reduce (Fattinger et al., 2017; Moreira et al., 2021; Ngo et al., 2013) or enhance SWA (Huwiler et al., 2022; Leach et al., 2024). These discrepancies may be attributed to differences in the applied stimulation protocols, with some favoring and some bypassing KCs. Viewing KCs as a probabilistic phenomenon, specific stimulation parameters related to *irregularity* and *novelty detection*, such as long ISIs, fast rise-and-fall times, and high stimulus intensities, all together modulate its probability of occurrence (Bastien & Campbell, 1992). Also, the phase within a sleep cycle (Halász, 2005) and other brain-state fluctuations (Dimitriades et al., 2024) may play a role. Studies reporting global SWA enhancement employed long stimulation breaks (OFF windows) of 8 to 10 seconds, significantly increasing the likelihood of evoking subsequent KCs. Such evoked KCs can also explain why, in these studies, the largest SWA increase was observed immediately after stimulation breaks, at the onset of ON windows (Huwiler et al., 2022; Krugliakova et al., 2022; Leach et al., 2024). In contrast, studies which observed a reduction in SWA with down-PTAS employed continuous stimulation protocols (Fattinger et al., 2017; Moreira et al., 2021) or implemented much shorter and regular stimulation breaks (Ngo et al., 2013). Such short ISIs may have led to sustained stimulus habituation, likely ideal for preventing evoked KC and allowing phase-specific effects to emerge.

The probabilistic nature of evoked KC may also explain the occasional observation of KCs following the second or third stimulus within a train (Davis et al., 1939; Roth et al., 1956), offering an explanation as to why the reduction of SWA following down-PTAS only started with the 5^th^ stimulus within the train. Given the shared directionality of the KC and phase-specific response for up-PTAS, pinpointing the specific stimulus within a train where the phase-specific outweighs the KC response is rather challenging (see **Fig. 1B**). It appears, however, that with each successive stimulus within a train, the probability of KC occurrence decreases, while phase-specific effects increasingly dominate.

Given the diverging electrophysiological response patterns following isolated stimuli and those presented consecutively within trains, it seems likely that the underlying neuronal mechanisms are also fundamentally distinct, akin to how different types of slow waves are thought to arise from different sources (Bernardi et al., 2018; Siclari et al., 2014). With this in mind, it seems possible that the KC and phase-specific response impact distinct sleep functions, with phase-specific responses particularly influencing functions associated with the targeted area (Fattinger et al., 2017; Sousouri et al., 2022). By disentangling PTAS responses, researchers may be able to establish causal links with behavioral outcomes and specific slow-wave subtypes, potentially represented by the distinct PTAS responses. Beyond a better understanding of basic sleep mechanisms, these insights could be essential for translating PTAS into clinical applications. Depending on the specific neuronal networks affected in a given clinical population, targeting one system over the other may lead to more effective outcomes—a distinction that could be particularly relevant in home or long-term settings (e.g., Kasties et al., 2024).

## Conclusion

Auditory stimuli are highly effective at evoking K-complexes (KCs). This property persists when stimuli are presented phase-locked to ongoing slow waves. Here, it was the *timing between stimuli* which determined whether a *phase-specific* or a *KC-mediated* response would unfold. These two distinct electrophysiological outcomes of PTAS, likely reflecting the engagement of different neural circuits associated with specific sleep-related functions, underscore the necessity of a nuanced mechanistic understanding of sleep neuromodulation, particularly PTAS. Such understanding is essential for establishing robust causal links between neural oscillations and behavior, advancing both fundamental neuroscience and clinical applications.

## Limitations

This study encompasses certain limitations that warrant acknowledgment. While the used datasets allowed the assessment of phase-specific effects following both up- and down-PTAS, the respective studies were originally not designed to investigate the impact of different ISIs. For instance, the employed ON|OFF window design, with a window duration of 6 s for up-PTAS, artificially interrupted continuous stimulus trains, resulting in borderline trial numbers for longer train lengths. This limitation prevented us from analyzing the responses to specific stimuli within a train, as doing so would have further reduced trial numbers. Additionally, stimuli were grouped post-hoc based on their stimulation-free periods prior to that stimulus. Consequently, the presentation of stimuli with shorter or longer stimulation-free periods was not experimentally controlled, potentially confounding this variable with other factors, such as variations in sleep depth. This is also why the interpretation of chosen ISI cutoffs (1 and 5 s) should be approached with caution. These cutoffs represent a trade-off between ensuring sufficient separation between stimuli to observe distinct responses and maintaining an adequate number of stimuli for robust statistical analysis. Lastly, the two studies recruited a rather small number of different individuals and differed in certain stimulation protocol settings, including different ON|OFF window durations, different detection electrode location, and different stimulation opportunity periods (see Methods). These differences most notably limited our ability to directly contrast responses between up- and down-PTAS. Although initial evidence suggests that the observed locality of the phase-specific response indeed depends on the location of the detection electrode (Fattinger et al., 2017), a systematic investigation of this relationship is needed to definitively resolve this question. Thus, while this study presents the very first compelling argument that ISIs are indeed a key factor for predicting PTAS responses, these findings should be validated in a study that is specifically designed to test this hypothesis.

## Acknowledgements

We are deeply grateful to the members of Prof. Reto Huber’s lab and the SleepLoop consortium for their invaluable constructive feedback and stimulating discussions. Their collaborative spirit significantly contributed to the development of this work. This research was supported by the Swiss National Science Foundation (SNF) under grant number 320030_179443 and is part of the HMZ Flagship grant *SleepLoop* under the umbrella of *Hochschulmedizin Zürich*, Switzerland.

## Methods

### Datasets

For the presented reanalysis, hd-EEG recordings (128 channels) from previously conducted studies were reanalyzed (Krugliakova et al., 2022; Leach et al., 2024; Sousouri et al., 2022). In the first dataset, N=18 participants underwent three nights: a night with up-PTAS, down-PTAS, and a night without stimulation as described in Sousouri et al. (2022). During the first 2.5 h of NREM sleep, 50 ms pink noise stimuli were presented (ISI *≥* 0.5 s) in an ON|OFF window design. Using that design, ON windows (6 s), allowing stimulation, took turns with OFF windows (6 s), withholding stimulation. The detection electrode, used for real-time PTAS during NREM sleep, was placed next to channel Fp2. This dataset was used previously (Krugliakova et al., 2022; Leach et al., 2024; Sousouri et al., 2022). For *this* reanalysis, we included hd-EEG data with and without up-PTAS from N=10 participants as reported in Sousouri et al. (2022) (mean±sd: 23.70±1.70 years old; all right-handed; 6 females, 4 males). Nights with down-PTAS were not considered due to an insufficient number of trials containing long stimulation trains. Instead, down-PTAS data were obtained from a second dataset, comprising N=14 participants (mean*±*sd = 23.25*±*2.53 years old; all right-handed; 10 females, 4 males) who underwent two nights: one night with and another without down-PTAS (Leach et al., 2024). The stimulation protocol was identical apart from longer ON (16 s) and OFF windows (8 s), as well as a longer stimulation opportunity (whole night), resulting in more trials with extended stimulus trains. The detection electrode was placed next to channel C3.

### Software

EEG analyses and statistics were performed in Matlab (R2022a & R2024b; The Math-Works, Inc., Natick, Massachusetts) implementing functions from the EEGLAB toolbox (v2023.1; Delorme & Makeig, 2004). Circular outcome measures (e.g., mean phase) were computed with functions from the toolbox for circular statistics (Berens, 2009).

### Preprocessing

Hd-EEG data was sleep-scored, artefact-corrected, and preprocessed as previously reported (Leach et al., 2024; Sousouri et al., 2022). To ensure consistent preprocessing across both datasets, filter settings were adapted to match those reported in Leach et al., 2024.

### Referencing

For time-frequency analyses, hd-EEG data was referenced to the mean of all channels located above the ear (up to 109 channels for up-PTAS; 110 channels for down-PTAS). Only those channels located above the ear were included in further analyses to minimize artifact contamination. For the phase estimation of auditory stimuli, hd-EEG data was referenced to linked mastoids.

### Time-frequency analysis

Time-frequency analyses for both up- and down-PTAS were performed as reported in Leach et al., 2024, separate for stimuli with short and long ISIs. To eliminate participant and frequency biases irrespective of the duration of the analyzed time window, spectral power values were normalized by dividing each value by the average spectral power value of both nights (sample- and frequency-wise). An automated outlier detection routine removed trials with clearly bad data. Specifically, trials where the maximum spectral power value across channels, samples, and frequencies, or the mean EEG signal amplitude across channels and samples exceeded six standard deviations from their respective mean across trials were excluded.

### Inter-stimulus interval (ISI)

Stimuli with long ISIs followed the previous stimulus with a distance of at least 5 seconds. Time-frequency responses of stimuli with long ISIs are depicted in **Fig. 1A** and **Fig. 2A**. Those with short ISIs followed the previous stimulus at a distance of 1 second or less. Whenever subsequent stimuli fulfilled this condition, they were considered to belong to the same train. For instance, a train of *five* stimuli consisted of *five* stimuli with an ISI of no longer than 1 second. The stimulation-free period prior to the first stimulus was not of interest. **Fig. 1B** and **Fig. 2B** depict the average response to a stimulus and all subsequent stimuli within the train rather than the response to a stimulus at a particular train position. With the assumption that a phase-specific response would persist with subsequent stimuli, such an approach increases trial numbers. The number of trains containing the respective number of stimuli is summarized in **Supplementary Fig. 2A**.

### Phase estimation of auditory stimuli

Phase estimations for stimuli of both up- and down-PTAS were computed as reported in Leach et al., 2024. Phase distributions are reported in **Supplementary Fig. 1**.

### Topographies

Absolute spectral power values were averaged across reported time- and frequency ranges, specific to the respective analysis.

### Statistics

For topographical comparisons between conditions (STIM−SHAM), paired Student’s *t*-tests (α = .05, two-tailed) were performed (channel-wise). To account for multiple comparisons, non-parametric cluster-based statistical mapping was applied. Effect sizes are reported by reporting the mean and maximum Hedge’s *g* value of significant channels.

### Non-parametric cluster-based statistical mapping

In short, the condition label (STIM or SHAM) was pseudorandomly assigned to participants, resulting in a random switch between conditions. The number of permutations is restricted by the number of participants (2^N^ − 1). Hence, the data was permuted 5000 times for down-PTAS and 1000 times for up-PTAS. Each permutation resulted in mutually exclusive condition labels. Paired Student’s *t*-tests were performed in each permutation, and the maximum cluster size of significant neighboring electrodes was computed separately for positive and negative *t*-values. This resulted in two distributions of cluster sizes, one for positive and one for negative *t*-values, with as many values as permutations performed. For topographical comparisons, the 97.5th percentile in each distribution was defined as the critical cluster size threshold. When the original data showed a cluster of significant electrodes equal to or larger than the critical cluster size threshold, this cluster of electrodes was considered significant. Multiple clusters could coexist and were reported as such when observed.

## Supplementary figures

**Supplementary Fig. 1:**
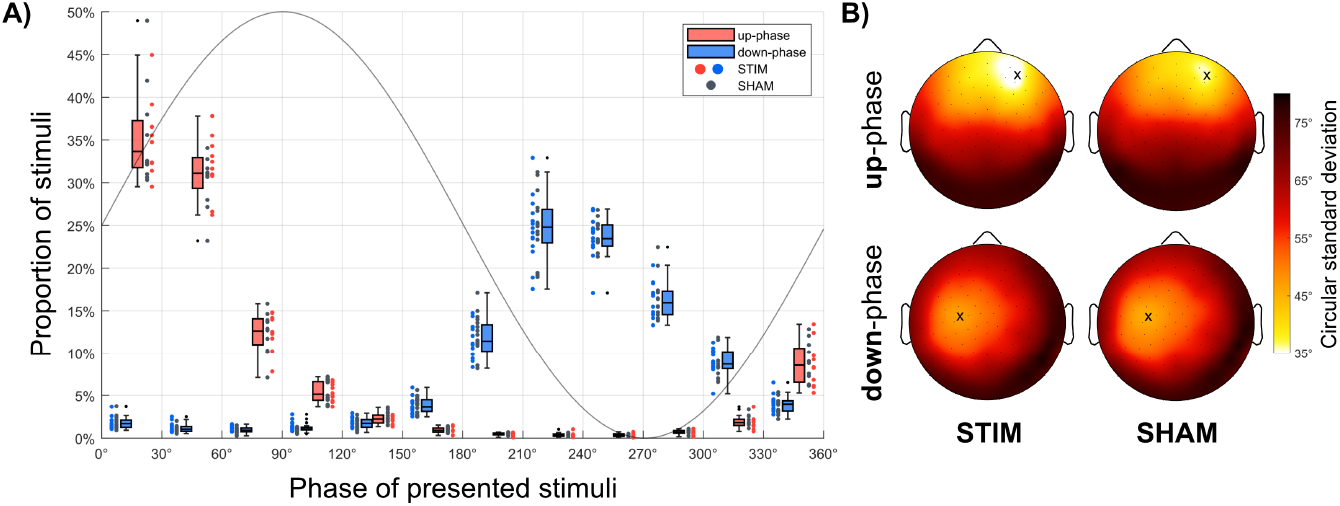
Phase precision of delivered stimuli. **(A)** Proportion of stimuli falling within specific phase bins (30° increments). The x-axis marks the boundaries of each phase bin. Proportions were calculated individually for each participant, and scatter-box plots show the distribution across participants. The color of the box plots reflects the targeted phase (red: up-phase targeting; blue: down-phase targeting), while the dot colors represent the stimulation condition (red and blue: STIM; grey: SHAM). **(B)** Topographic phase precision, expressed as the circular standard deviation of phase values at the times of stimulation. For each electrode, the circular standard deviation was computed per participant, then averaged across participants. Topographic maps of phase precision are shown separately for up- and down-phase targeting, as well as for STIM and SHAM nights. The black cross marks the detection electrode, where phase precision is highest.

**Supplementary Fig. 2:**
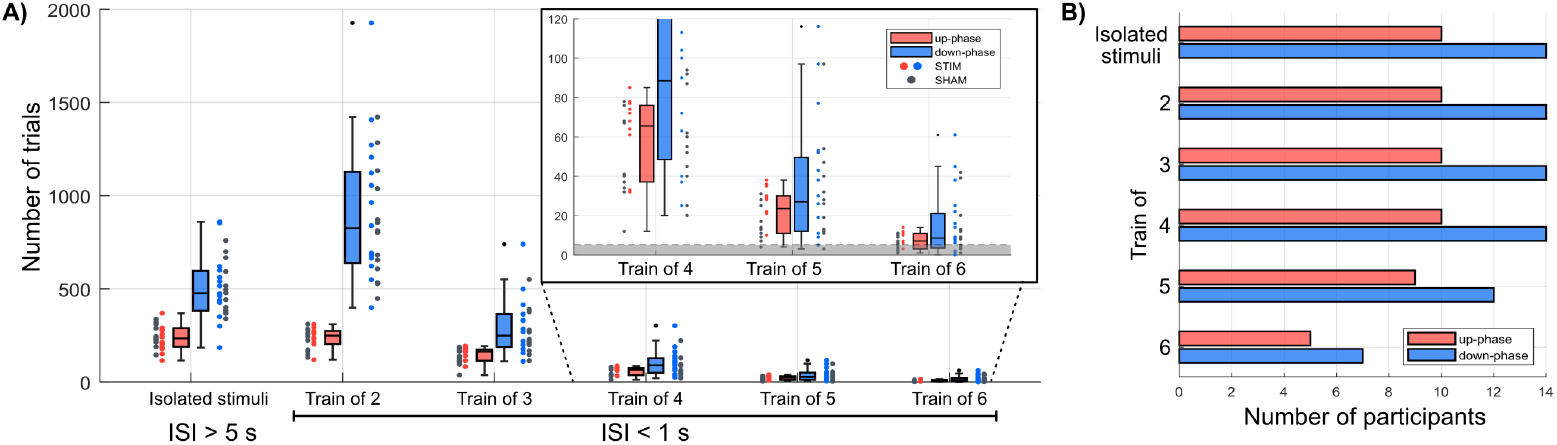
Number of trials. **(A)** Scatter-box plots illustrate the number of isolated stimuli per night, defined as stimuli following a stimulation-free period of *≥* 5 s, alongside the number of stimulus trains of varying lengths (2–6 consecutive stimuli) with an inter-stimulus interval (ISI) of ≤ 1 s. The availability of trials diminished with increasing train length. Number of trials is presented separately for nights with up-phase-targeted (red) and down-phase-targeted (blue) auditory stimulation (PTAS). Notably, the number of trials was generally higher in nights with down-compared to up-PTAS, attributed to longer ON windows (16 s vs. 6 s), higher ON/OFF window ratios (16/8 vs. 6/6), and longer overall stimulation periods (entire night vs. first 2.5 hours). Colored dots indicate the number of trials per stimulation night, while grey dots represent SHAM nights. The inset plot in the top right provides a zoomed-in view of the number of trains for the longest stimulus trains. The grey shaded area indicates participants with less than 5 trials per night. **(B)** Bar plot displaying the number of participants with *≥* 5 trials per night for trains with varying numbers of stimuli. The longest train length which was analyzed was 5 consecutive stimuli.

**Supplementary Fig. 3:**
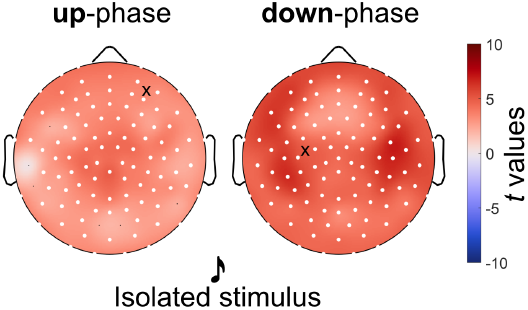
Theta activity following isolated stimuli. Topography of absolute theta activity (4 – 8 Hz; 300 – 700 ms following stimulus onset) in response to isolated stimuli, that is, stimuli following a stimulation-free period of *≥* 5 s. Responses are displayed either for stimuli targeting the up-phase (left) or down-phase (right) of slow waves. **Symbols:** The black cross indicates the detection electrode. White dots represent electrodes with significant differences between conditions (STIM vs. SHAM, paired *t*-test, cluster corrected).

**Supplementary Fig. 4:**
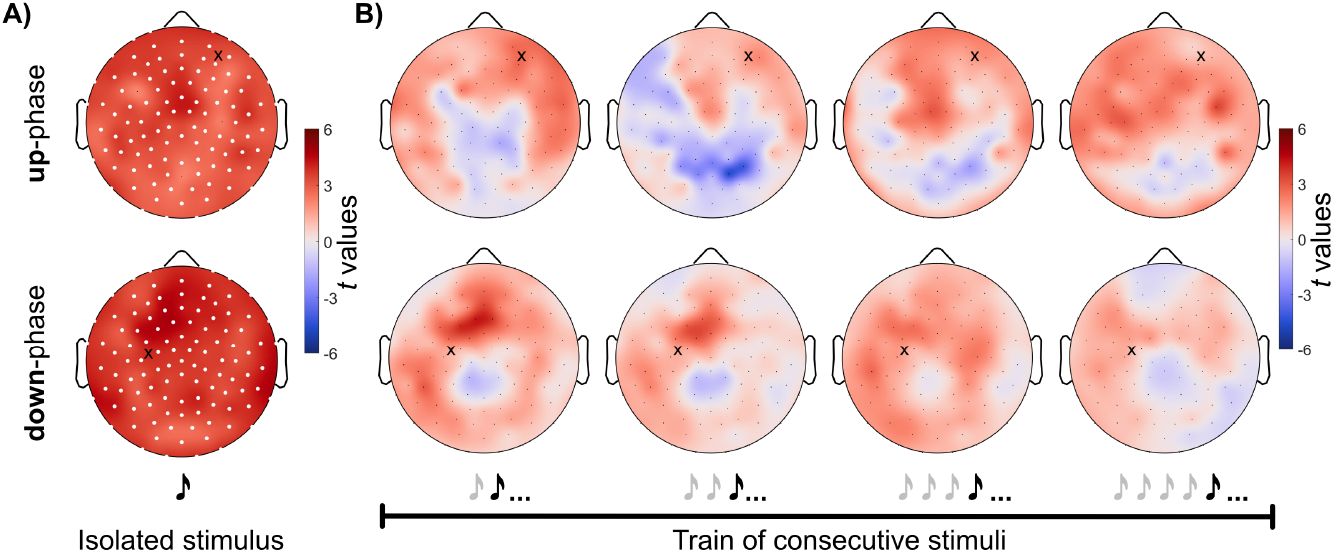
Sigma activity response over time. **(A)** Topography of absolute sigma activity (12 – 16 Hz; 0.9 – 1.5 s following stimulus onset) in response to isolated stimuli, that is, stimuli following a stimulation-free period of *≥* 5 s. Responses are displayed either for stimuli targeting the up-phase (top) or down-phase (bottom) of slow waves. **(B)** Topography of absolute sigma activity (0.9 – 1.5 s) in response to stimuli within a train of consecutive stimuli, that is, stimuli with an inter-stimulus interval (ISI) ≤ 1 s. The average response to a stimulus and all subsequent stimuli within the train rather than the response to a stimulus at a particular train position is shown (indicated by the ellipsis, the three dots). Note that the sigma response is only present following isolated stimuli. **Symbols:** The black cross indicates the detection electrode. White dots represent electrodes with significant differences between conditions (STIM vs. SHAM, paired *t*-test, cluster corrected).

